# Amazon Mechanical Turk as a platform for borderline personality disorder research

**DOI:** 10.1101/316356

**Keywords:** MTurk, crowdsourcing, borderline personality disorder, remission, recovery

## Abstract

Researchers investigating the psychological processes underlying specific mental health problems often have difficulties achieving large enough samples for adequately powered studies. This can be particularly problematic when studying psychopathology with low base rates in typical samples (i.e., undergraduate and community). A relatively new approach to recruitment and testing employs online crowdsourcing to rapidly measure the characteristics and behavior of large numbers of people. We tested the feasibility of researching borderline personality disorder (BPD) in this manner using one large crowdsourcing site, Amazon Mechanical Turk (MTurk). Specifically, we examined prevalence rates of psychopathology in a large MTurk sample, as well as the demographic, psychosocial, and psychiatric characteristics of individuals who met criteria for BPD. These characteristics were compared across three groups: those who met criteria for BPD currently, those who met criteria for remitted BPD, and those who had never met criteria for BPD. The results suggest that MTurk may be ideally suited for studying individuals with a wide range of pathology, from healthy to intensely symptomatic to remitted.

## Introduction

The present study examines the feasibility of using Amazon Mechanical Turk (MTurk), an on-line data collection platform, to study borderline personality disorder (BPD). Specifically, we examine, across two studies, the correlates of BPD. We seek to recreate in data collected from MTurk participants the nomological network built around BPD in the literature, as well as prevalence rates established by current epidemiological research. We examine demographic, psychosocial, medical, and basic personality characteristics among three subgroups within our sample: those who meet criteria for current BPD, those who meet criteria for remitted BPD, and those who do not meet criteria for either. We evaluate the appropriateness of MTurk as a venue for studying BPD in the context of these findings.

### MTurk

MTurk is an online labor market where “workers” choose and complete small tasks, referred to as human intelligence tasks (HITs). Each task is listed with a description of the activity, time to complete, and proposed payment for completion, and workers are paid through the website after work is submitted and checked. HITs can be completed on a personal computer from anywhere.

There are several crowdsourcing markets, of which MTurk is currently the largest and most studied non-probability sample available to researchers (Chandler & Shapiro, 2016). MTurk is increasingly used in psychological research because it facilitates rapid collection of large datasets at relatively low cost (Chandler & Shapiro, 2016). Also, workers can be selected based on specific qualifications (e.g., native English-speakers, workers in the U.S., workers with high ratings of completion of other HITs), or pre-screened using a brief initial HIT to sort subjects by other characteristics of interest prior to enrollment in a larger study.

MTurk is also attractive to researchers because it provides access to individuals who would not normally be represented in a clinic-based or convenience sample (Gosling & Mason, 2015). This group may include people with significant symptoms who have never presented for clinical care, and also those whose geographic locations, schedules, or preferences would dissuade them from presenting as research subjects. Demographics of MTurk workers are diverse, though not perfectly representative: workers are younger and better educated than the general population, and are primarily European or Asian American (Paolacci & Chandler, 2014). They have lower incomes and are more likely to be un- or under-employed (Corrigan, Bink, Fokuo, & Schmidt, 2015). Nonetheless, MTurk samples better match community samples than do typical research populations (e.g., college students and community volunteers; (Berinsky, Huber, & Lenz, 2012; Chandler & Shapiro, 2016).

Mental health problems are also well represented on MTurk as workers have anxiety and depression at levels comparable to the general population (Shapiro, Chandler, & Mueller, 2013). Additional research has examined personality disorders among MTurk workers, although this work has focused largely on narcissistic personality disorder. In fact, while the vast majority of research has examined NPD, only one of 98 studies examining personality disorders in MTurk published in two major psychology journals in the past three years focused on borderline personality disorder (Miller, Crowe, Weiss, Maples-Keller, & Lynam, 2017).

There are some limitations of the MTurk sample worth noting (Miller, Crowe, Weiss, Maples-Keller, & Lynam, 2017). Participants may be non-naïve, having potentially completed numerous psychological tasks on MTurk. They may be familiar with either the tasks or the methods used for checking attention and response validity. Also, the MTurk participant pool may be smaller than the apparent millions of workers suggested by the platform: some data suggest that a small percentage of workers complete a substantial proportion of the HITs (Chandler, Mueller, & Paolacci, 2014). This requires researchers who hope to replicate or validate findings in repeated MTurk studies to attend closely to worker IDs in order to avoid recruiting duplicate individuals in subsequent samples.

### BPD characteristics

BPD is a disorder that is characterized by serious pervasive dysfunction and instability in personal relationships, affect, perceptual experience, and self-image. Impulsivity and intentional self-harm are common. BPD symptoms are associated with negative relationship experiences such as chronic relationship stress, partner dissatisfaction, relationship conflict, abuse, and unwanted pregnancy. In addition, adults with BPD report severe difficulty with multiple aspects of parenting, including the transition to parenthood, infant care, and early attachment (Zanarini, Frankenburg, Reich, Wedig, et al., 2015). Finally, given the substantial distress and impairment associated with this disorder, it is not surprising that it is associated with substantial health care utilization costs (Soeteman, Hakkaart-van Roijen, Verheul, & Busschbach, 2008).

## BPD’s Nomological Network

### Prevalence and Comorbidities

Data collected from a community sample of over 34,000 individuals in the National Epidemiologic Survey on Alcohol and Related Conditions (NESARC) found that 5.9% of adults in the United states meet diagnostic criteria for BPD (Hasin & Grant, 2015). The NESARC found more frequent BPD among people who were female, low income, younger than 30, separated or divorced, Native American or African American. Others have examined co-morbidities in BPD: anxiety disorders, substance use disorders, and other personality disorders were most frequent (Swartz, Blazer, George, & Winfield, 1990; Tomko, Trull, Wood, & Sher, 2014). Lifetime prevalence of comorbid mood and trauma related disorders is 85%, prevalence of comorbid substance use disorders is 78% (Tomko, Trull, Wood, & Sher, 2014), and prevalence of comorbid chronic pain is 30% (Heath, Paris, Laporte, & Gill, 2017). BPD also increases risk for other medical problems (reviewed and discussed in (Iacovino, Powers, & Oltmanns, 2014).

### Psychiatric medication use

Zanarini and colleagues (Zanarini, Frankenburg, Reich, Harned, & Fitzmaurice, 2015) investigated psychiatric medication use in BPD compared to other personality disorders. They found that people with BPD took more antidepressants (1.3 times more), antipsychotics (2.6 times more) and mood stabilizers (3 times more) than did comparison subjects. Prescription opiate use was also more common in BPD than in other personality disorders (Frankenburg, Fitzmaurice, & Zanarini, 2014).

### Self-harm

Among inpatients with BPD, 90% endorse history of self-injury and more than 70% endorse history of suicide attempt(s) (Goodman et al., 2017). Other studies in adult samples have also found high rates of suicidal ideation and attempts among people with BPD (Wedig, Frankenburg, Reich, Fitzmaurice, & Zanarini, 2013; Wedig et al., 2012) (Paris & Zweig-Frank, 2001) (Boisseau et al., 2013) (Pompili, Girardi, Ruberto, & Tatarelli, 2005). Between 5-10% complete suicide (Black, Blum, Pfohl, & Hale, 2004; Paris, 2006) (Pompili et al., 2005).

### Basic personality traits

Research has examined personality disorders as clusters of extreme variants of basic personality traits in the five factor model (FFM) (e.g., (Trull & Widiger, 2013) -- a classification system of normative personality traits that fall into five broad domains: neuroticism, extraversion, openness to experience, agreeableness, and conscientiousness. In a recent meta-analysis, Samuel & Widiger (2008) found that BPD bears strong positive relations to neuroticism (*r* = .54) and negative relations to agreeableness (*r* = -.24) and conscientiousness (*r* = -.29).

These correlates are consistent with the FFM facet profile for BPD generated by experts (Lynam & Widiger, 2001). Similarly, (Hopwood & Zanarini, 2010) found that FFM neuroticism was significantly correlated to the affective (*r* = .50), cognitive (*r* = .39), and interpersonal domains (*r* = .45) of BPD. In addition, FFM agreeableness was inversely correlated to impulsive symptoms (*r* = -.23), and agreeableness and extraversion were inversely correlated to BPD symptoms overall.

### Remission and recovery in BPD

Research suggests that BPD symptoms remit over time for most people (75-99%) within a few years, and 40-60% achieve functional recovery (Zanarini, Frankenburg, Reich, & Fitzmaurice, 2012). Although these numbers are promising, people in remission from BPD are much less likely to exhibit high levels of functioning socially or in the community. For example, while people recovered from BPD are significantly more likely than non-recovered individuals with BPD to have married or lived with a romantic partner, these same individuals are more likely to have given up or lost custody of a child (Zanarini, Frankenburg, Reich, Wedig, et al., 2015). These findings highlight the complexity of what it means to be in remission versus recovered and suggests that more research is needed to examine quality of life among these diagnostic groups.

### Current Study

The current study seeks to replicate, in an MTurk sample, the nomological network that surrounds BPD more broadly. We recruited more than 3,000 people via MTurk to answer questions about demographics, emotions, and behavior. Self-report surveys were administered including instruments to measure BPD symptoms now and in the past, as well as basic personality traits. A subset of this large group (~ 700 people) went on to a second experiment, which included measures of BPD, emotion regulation, impulsivity, depression and anxiety. We report on demographics and symptoms in people with current BPD, remitted BPD, and no history of BPD.

## Methods

### Ethics

This protocol and consent materials were approved by the [relevant institutional – specifics removed for masked file] Review Board. All participants were presented with the consent form online at the start of both parts one and two of the study. The consent information described anticipated risk (minimal risk due to potential discomfort when answering personal questions), confidentiality, a reminder that agreement to participate is voluntary and could be revoked, and contact information for the study team.

### Participants

Recruitment on MTurk was restricted to workers in the United States. Workers were invited to complete a 15-25 minute HIT about their “life, emotional state, and behavior” for $0.40 compensation (HIT1). The study description also stated that some HIT1 participants would be invited to participate in a subsequent, more highly paid, HIT (HIT2 – see below). The MTurk site was linked to a survey platform on Qualtrics, where participants completed the surveys. Participants provided their MTurk ID and a survey completion code from Qualtrics to link their MTurk profile to their Qualtrics survey responses.

The initial group of HIT1 completers totaled 3,633 participants. We excluded participants for completing the task too quickly (< 9 minutes), for missing any of the three embedded attention check items, or for extreme frequency in responding (i.e. those who chose the same response choice at statistically outlying rates). After these exclusions, 3,132 participants remained. We also excluded people who reported color blindness, learning disability, traumatic brain injury, or schizophrenia. The ability to correctly perceive colors was important to a social cognition task included in HIT2, but not reported here. A final sample of 3,021 participants was used for analysis of HIT1 scales.

We invited a subset of HIT1 participants to participate in a second set of surveys and tasks, “HIT2”. We sorted the HIT1 completers into ten groups by gender (M/F) and BPD symptom level (based on SCID II questionnaire, endorsed currently having 0-2, 3-4, 5-9, 10+ symptoms, or having <5 current symptoms, but ≥ 5 past symptoms). For this process, we defined current as within the past 2 years. HIT1 and HIT2 ran simultaneously: as people completed HIT2, we aimed to fill each of the 10 categories with 30 participants. When each category was full, we stopped inviting HIT1 participants for that category. This approach allowed us to oversample for BPD symptoms and for male participants in HIT2, but it does result in a HIT2 sample that is not representative of the general MTurk population. This process resulted in an initial sample of 1,134 participants, all of whom had completed HIT1. These categories were used only for recruitment. Please see below in the Methods section regarding the SCID-II-PQBPD scale for the definitions of the three groups we defined for analysis.

HIT2 participants experienced a consent process as described for HIT1, however it described a longer HIT of ~ 40 minutes, including social games, and compensation of $2.00 plus a possible $2.00 bonus for excellent game performance. We excluded people who were ultimately excluded from HIT1 analyses (due to extreme responses, etc.; n = 243). We then excluded people who did not pass HIT2 attention checks (n = 52), or missed some survey questions (n = 128). A final sample of 711 participants was used for analysis of HIT2 surveys.

## HIT1 Measures

### Demographics and medical history questionnaire

Participants responded to a series of questions asking about demographic characteristics, past medical and psychiatric history, and current medications. Medications were manually read and divided into categories by psychiatrists (SKF and ER).

### International Personality Item Pool Representation of the NEO PI-R Short Form (IPIP-NEO SF)

The short form of the IPIP-NEO (Maples-Keller, et al., 2017) is comprised of 60 items that measure the thirty facets of the five domains (neuroticism, openness, conscientiousness, agreeableness, and extraversion) of the Five Factor Model (FFM). Twelve items comprise each domain, with two items representing each of the thirty FFM facets. In the present study, alpha coefficients for the domain scales ranged from .57 to .77.

### Structured Clinical Interview for DSM-IV Axis II Personality Disorders Self-report Questionnaire (SCID-II-PQ-BPD)

We employed the 15 BPD questions from the SCID-II personality disorders self-report questionnaire, which assess the nine diagnostic criteria for BPD as outlined in DSM-IV (First, Gibbon, Spitzer, Williams, & Benjamin, 1997;First, Spitzer, Gibbon, Williams, Benjamin, 1994). For each item that asked participants if they had experienced a given symptom, they were given four response choices: “Yes, that happened this month”; “Not this month, but in the last 2 years”; “Not in the last 2 years, but in the past”; and “No.” The alpha coefficient for the SCID-II-PQ-BPD in the current study was .88.

### Participant Group Assignments for Analysis

We used the SCID-II-PQ-BPD responses to define three groups. First, the fifteen questions were used to code binary responses to each of the nine DSM criteria. If participants reported at least five of the nine DSM criteria in the last two years (we combined responses “yes, that happened this month” and “not this month, but in the last 2 years”), they were assigned to the group “Current BPD” (n = 1,020). If participants reported at least five of the nine criteria in their lifetime, but not within the last two years, they were assigned to the “Remitted BPD” group (n = 865). The remaining participants were assigned to the “Never BPD” group (n = 1,136).

Participants in HIT1 also completed several tasks that are not discussed in this paper. These were measures of psychopathy, narcissism, autism, a brief writing sample in response to the prompt “Please tell us about yourself,” and a five-round Trust Game.

## HIT2 Measures

### Beck Anxiety Inventory (BAI)

The BAI is a 21-item self-report measure assessing various symptoms of anxiety (Beck, Epstein, Brown, & Steer, 1988; Fydrich, Dowdall, & Chambless, 1992). Participants rate the symptoms (e.g., “Numbness or tingling”) on a scale from 0 (Not at all) to 3 (Severely - it bothered me a lot) indicating how much he/she has been bothered by that symptom during the past month, with a total possible score of 63 points. The BAI demonstrated good reliability in the present study (α = .94).

### Borderline Symptom List 23 (BSL-23)

The BSL-23 consists of 23 items assessing BPD symptoms experienced in the past week (Bohus et al., 2009). Participants indicate how the extent to which they experienced each item on a scale from 0 (Not at all) to 4 (Very strong). Sample items include “I was lonely” and “I suffered from shame.” In the current study, the BSL-23 had a coefficient alpha of .95.

### World Health Organization Quality of Life - BREF (WHOQOL-BREF)

The WHOQOL-BREF (The WHOQOL Group, 1998) is a 26-item scale designed to assess perceptions of physical health, psychological health, social relationships, and environment. Item responses are given on a five-point Likert scale. Reliability for this scale was .93.

### Beck Depression Inventory Second Edition (BDI-II)

The BDI-II (Beck, Steer, & Brown, 1996) consists of 21 items and is used to assess the existence and severity of depression symptoms in the past two weeks as outlined in the DSM-IV. There is a four-point scale for each item ranging from 0 to 3, with higher scores indicating elevated symptom levels. The BDI demonstrated strong reliability in the present study (α = .95).

### Barratt Impulsiveness Scale (BIS-11)

The BIS-11 (Patton, Stanford, & Barratt, 1995) was designed to assess multiple facets of impulsivity. The BIS-11 comprises 30 items, all on a 4-point scale ranging from “Rarely/Never” to “Almost Always/Always.” Reliability for this was measure was .86 overall and .65-.76 for the subscales in the current study.

### Difficulties in Emotion Regulation Scale (DERS)

The DERS (Gratz & Roemer, 2004) is a 36-item measure that assesses a person’s beliefs and feelings when he/she becomes upset, as well as a person’s perceptions of how they experience emotion. Items are measured on a five-point Likert scale. Coefficient alpha was 0.87.

### Positive and Negative Affect Scale (PANAS)

The PANAS (Watson, Clark, & Tellegen, 1988) comprises two mood scales, one assessing positive affect and the other assessing negative affect. There are 20 items in the measure, each on a five-point Likert scale. Both the positive and negative affect scales demonstrated good reliability (α = .92 and α = .89, respectively).

After responding to these self report measures, HIT2 participants also completed two tasks not discussed here: one 10 round Cyberball game (Williams & Jarvis, 2006; Williams, Yaeager, Cheung, & Choi, 2012) followed by an associative learning task focused on social cognition.

## Results

### Characteristics of the overall sample

The demographics of our sample (HIT1) were consistent with past research on the MTurk population (Berinsky, Huber, & Lenz, 2012; Shapiro, 2013; Buhrmester, Kwang, & Gosling, 2011; Paolacci &Chandler, 2014). Participants were overwhelmingly female (66.1%), white (74%), middle-aged (*M* = 37.12 years, range 18 – 81, *SD* = 12.26), heterosexual (84.9%), and well educated (86.2% had completed at least some college). Only a small fraction (20.4%) were currently in school, but most (70.3%) were employed at least 10-20 hours/week, and 43.2% were employed full-time.

### Demographics of people with current, remitted, and never BPD

We found that 1020 HIT1 participants (33.8%) met DSM criteria for BPD in the last 2 years – we termed this group “**Current BPD**”. Another 28.6% of HIT1 participants endorsed having had 5+ DSM criteria for BPD sometime during their life, but having ≤ 5 DSM BPD criteria in the past two years. This group was termed “**Remitted BPD**.” The remaining participants, who had fewer than five lifetime BPD symptoms, belong to the “**Never BPD**” group. Of note, only 1% of the overall sample, and 2.2% of the Current BPD group had been previously <u>diagnosed</u> with BPD (**Table 1**).

**Table 1.**
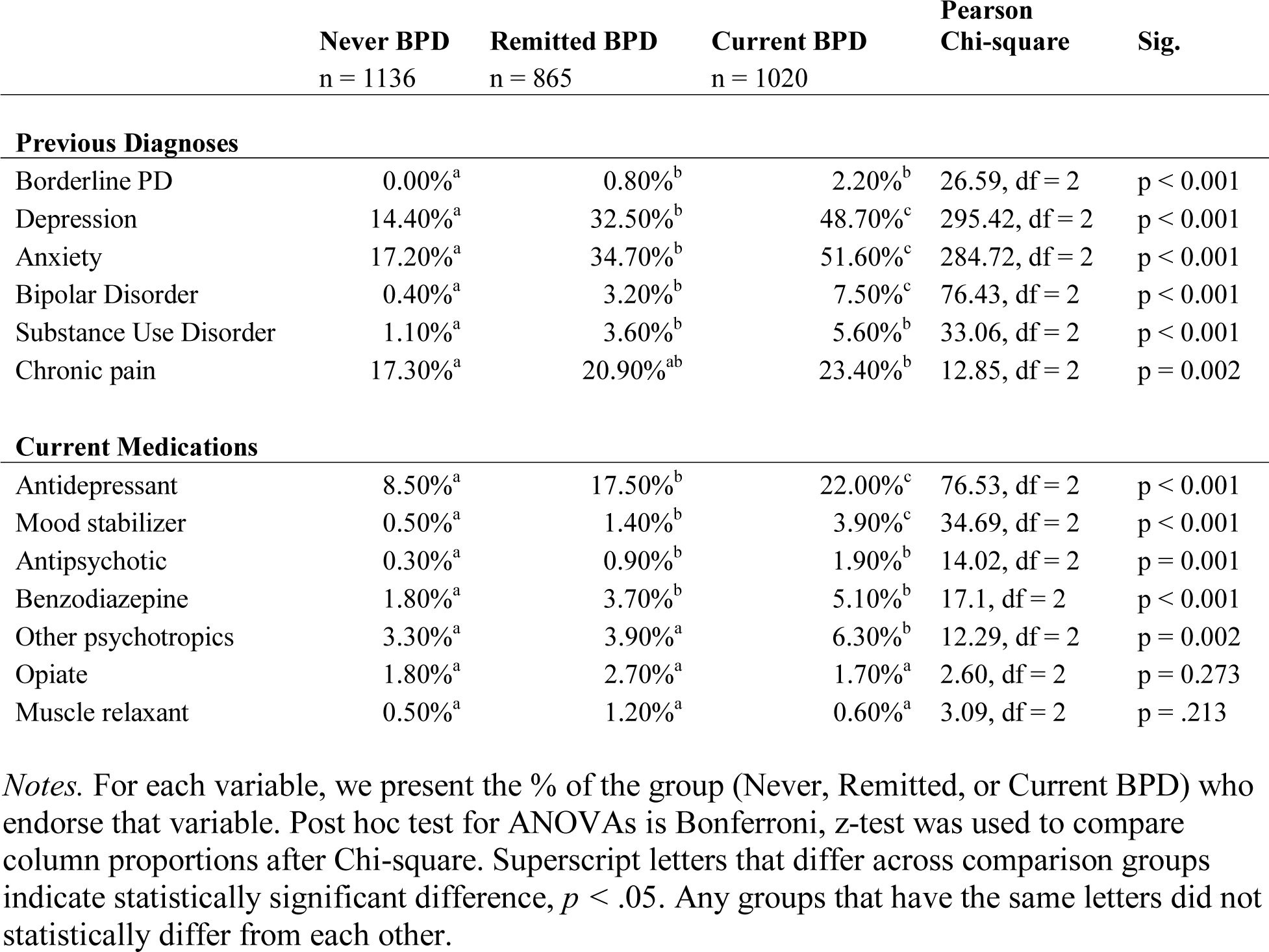
Previous diagnoses and medications in HIT1 sample (N = 3,021).

|  | **Never BPD**<br>n = 1136 | **Remitted BPD**<br>n = 865 | **Current BPD**<br>n = 1020 | **Pearson**<br>**Chi-square** | **Sig.** |
| --- | --- | --- | --- | --- | --- |
| **Previous Diagnoses** |  |  |  |  |  |
| Borderline PD | 0.00%<sup>a</sup> | 0.80%<sup>b</sup> | 2.20%<sup>b</sup> | 26.59, df = 2 | p < 0.001 |
| Depression | 14.40%<sup>a</sup> | 32.50%<sup>b</sup> | 48.70%<sup>c</sup> | 295.42, df = 2 | p < 0.001 |
| Anxiety | 17.20%<sup>a</sup> | 34.70%<sup>b</sup> | 51.60%<sup>c</sup> | 284.72, df = 2 | p < 0.001 |
| Bipolar Disorder | 0.40%<sup>a</sup> | 3.20%<sup>b</sup> | 7.50%<sup>c</sup> | 76.43, df = 2 | p < 0.001 |
| Substance Use Disorder | 1.10%<sup>a</sup> | 3.60%<sup>b</sup> | 5.60%<sup>b</sup> | 33.06, df = 2 | p < 0.001 |
| Chronic pain | 17.30%<sup>a</sup> | 20.90%<sup>ab</sup> | 23.40%<sup>b</sup> | 12.85, df = 2 | p = 0.002 |
| **Current Medications** |  |  |  |  |  |
| Antidepressant | 8.50%<sup>a</sup> | 17.50%<sup>b</sup> | 22.00%<sup>c</sup> | 76.53, df = 2 | p < 0.001 |
| Mood stabilizer | 0.50%<sup>a</sup> | 1.40%<sup>b</sup> | 3.90%<sup>c</sup> | 34.69, df = 2 | p < 0.001 |
| Antipsychotic | 0.30%<sup>a</sup> | 0.90%<sup>b</sup> | 1.90%<sup>b</sup> | 14.02, df = 2 | p = 0.001 |
| Benzodiazepine | 1.80%<sup>a</sup> | 3.70%<sup>b</sup> | 5.10%<sup>b</sup> | 17.1, df = 2 | p < 0.001 |
| Other psychotropics | 3.30%<sup>a</sup> | 3.90%<sup>a</sup> | 6.30%<sup>b</sup> | 12.29, df = 2 | p = 0.002 |
| Opiate | 1.80%<sup>a</sup> | 2.70%<sup>a</sup> | 1.70%<sup>a</sup> | 2.60, df = 2 | p = 0.273 |
| Muscle relaxant | 0.50%<sup>a</sup> | 1.20%<sup>a</sup> | 0.60%<sup>a</sup> | 3.09, df = 2 | p = .213 |

*Notes.* For each variable, we present the % of the group (Never, Remitted, or Current BPD) who endorse that variable. Post hoc test for ANOVAs is Bonferroni, z-test was used to compare column proportions after Chi-square. Superscript letters that differ across comparison groups indicate statistically significant difference, $p < .05$ . Any groups that have the same letters did not statistically differ from each other.

We observed some demographic differences among the three groups (**Table 1**). People with BPD were younger (Never BPD *M* = 41.03, *SD* = 13.48; Remitted BPD *M* = 37.87, *SD* = 11.67; Current BPD *M* = 32.17, *SD* = 9.27, *F*(2, 2989) = 155.70, *p* < 0.001, post hoc for all 3 comparisons p < 0.001). There were no significant group differences by gender (Never BPD = 66.6% female, Remitted BPD = 63.4% female, Current BPD = 67.9% female, (X^2^ (2) = 4.49, *p* = 0.11).

### Co-morbidities of people with current, remitted, and never BPD

We examined common co-morbidities in people with BPD (**Table 1**). As expected, people with BPD were more likely to endorse anxiety (X^2^ (2) = 284.72, *p* < 0.001, Cramer’s V = 0.307), depression (X^2^ (2) = 295.42, *p* < 0.001, Cramer’s V = 0.313), and bipolar disorder (X^2^ (2) = 76.43, *p* < 0.001, Cramer’s V = 0.159). For anxiety, depression, and bipolar disorder, the Remitted BPD represented an intermediate phenotype (post-hoc tests for all 3 comparisons significant at *p* < 0.05).

Participants with BPD were also more likely to endorse substance use disorders (X^2^ (2) = 33.06, *p* < 0.001, Cramer’s V = 0.105). Of note, in the overall sample, 3-4% of participants reported a diagnosis of substance use disorder (SUD), which is comparable to previous work on MTurk, where 4.3% endorsed having sought treatment for SUD (though 37.1% endorsed SUD-consistent symptoms) (Shapiro, Chandler, & Mueller, 2013). We did not use self-reports to screen for SUD symptoms here, we only asked about <u>diagnosis</u>. Here, the remitted group reported significantly more frequent SUD diagnosis than Never BPD participants (*p* < 0.05), and this was not significantly different than Current BPD group frequency.

Chronic pain was also more frequent in Current BPD than in Never BPD (23.4% versus 17.3%, X^2^ (2), *p* < .01, Cramer’s V = 0.065), however frequency in Remitted BPD was not significantly different than either of the other two groups.

People with BPD were more likely to be taking psychotropic medications, including antidepressants (X^2^ (2) = 76.53, *p* < 0.001, Cramer’s V = 0.159), mood stabilizers (X^2^ (2) = 34.69, *p* < 0.001, Cramer’s V = 0.107), antipsychotics (X^2^ (2) = 14.02, *p* < 0.001, Cramer’s V = 0.068), benzodiazepines (X^2^ (2) = 17.1, *p* < 0.001, Cramer’s V = 0.075), and other miscellaneous psychotropics (X^2^ (2) = 12.29, *p* < .001, Cramer’s V = 0.064). For anti-depressants and mood-stabilizers, frequency is at an intermediate level in the Remitted BPD group (significant differences between each of the 3 groups, *p* < 0.05). We did not observe significant differences in opiate or muscle relaxant use between groups.

### Relationship status of people with current, remitted, and never BPD

We found significant differences in both current relationship status (X^2^ (2) = 137.33, *p* < 0.001, Cramer’s V = 0.151) and parental status (X^2^ (2) = 50.31, *p* < 0.001, Cramer’s V = 0.129) in our sample (**Table 2**). Participants in the Current BPD group were less likely to be married (51.8% of Never BPD, 32.5% of Current BPD), and more likely to be in a relationship but not married (16.2% of Never BPD, 28.7% of Current BPD) or single (21.3% of Never BPD, 30.6% of Current BPD). Participants in the Remitted BPD group demonstrated an intermediate phenotype for frequency of participants married (46.4%) and in a relationship, not married (23.5%). Likelihood of being a parent decreases from Never BPD to Remitted BPD to Current BPD.

**Table 2.** Relationship status of full HIT1 sample (N = 3,021).

|  | **N: Never**<br>**BPD (%**<br>**of group)** | **R: Remitted**<br>**BPD (% of**<br>**group)** | **B: Current**<br>**BPD (% of**<br>**group)** | **Pearson chi**<br>**square** | **Sig.** |
| --- | --- | --- | --- | --- | --- |
| **Current Relationship**<br>**Status** |  |  |  | 137.328, df =<br>12 | p < 0.001 |
| Married | 51.80%<sup>a</sup> | 46.40%<sup>b</sup> | 32.50%<sup>c</sup> |  |  |
| In a relationship, not<br>married | 16.20%<sup>a</sup> | 23.50%<sup>b</sup> | 28.70%<sup>c</sup> |  |  |
| Divorced | 6.80%<sup>a</sup> | 4.70%<sup>b</sup> | 4.10%<sup>b</sup> |  |  |
| Separated | 1.10%<sup>a</sup> | 0.70%<sup>a</sup> | 1.30%<sup>a</sup> |  |  |
| Single | 21.30%<sup>a</sup> | 22.40%<sup>a</sup> | 30.60%<sup>b</sup> |  |  |
| Widowed | 1.70%<sup>a</sup> | 0.30%<sup>b</sup> | 0.30%<sup>b</sup> |  |  |
| Other | 1.20%<sup>a</sup> | 2.00%<sup>a</sup> | 2.50%<sup>a</sup> |  |  |
| **Parent** | 58.50%<sup>a</sup> | 52.80%<sup>b</sup> | 43.30%<sup>c</sup> | 50.313, df = 2 | p < 0.001 |

*Note.* Post hoc test for ANOVAs is Bonferroni, z-test was used to compare column proportions after Chi-square. Superscript letters that differ across comparison groups indicate statistically significant difference, $p < .05$ . Any groups that have the same letters did not statistically differ from each other.

### BPD symptoms endorsed in HIT1 and HIT2 surveys

We also examined the number and specifics of the BPD symptoms endorsed on the HIT1 assessment (SCID-II self-report questionnaire) (**Table 3**). People in the Current BPD group endorsed an average of 6.57 current symptoms and 1.85 remitted symptoms. People in the Remitted BPD endorsed an average of 2.78 current symptoms and 3.69 remitted symptoms. People in the Never BPD group endorsed 1.19 current symptoms and 1.19 remitted symptoms.

**Table 3.** BPD symptoms across diagnostic subgroups. Data presented as mean value, standard deviation.

|  | Never BPD | Remitted BPD | Current BPD | One-way ANOVA | Sig. |
| --- | --- | --- | --- | --- | --- |
| **Current DSM BPD symptoms (past 2 years)** | 1.19, 1.09<sup>a</sup> | 2.78, 1.19<sup>b</sup> | 6.57, 1.32<sup>c</sup> | F = 5592.21, df = 2 | p < 0.001 |
| **Remitted DSM BPD symptoms** | 1.19, 1.10<sup>a</sup> | 3.69, 1.62<sup>b</sup> | 1.85, 1.39<sup>c</sup> | F = 859.51, df = 2 | p < 0.001 |
| **HIT1 to HIT2 reliability** | **Time elapsed** |  |  | **Pearson correlation** | **Sig.** |
| **DSM-BPD sx to BSL23 score** | 16.80 days, 24.42 |  |  | 0.582 | p < 0.001 |
|  | Never BPD<br>n = 255 | Remitted BPD<br>n = 180 | Current BPD<br>n = 276 | One-way ANOVA | Sig. |
| **HIT2 scales** |  |  |  |  |  |
| **BPD symptoms** | 7.81, 11.01<sup>a</sup> | 14.93, 12.7<sup>b</sup> | 28.55, 8.53<sup>c</sup> | F = 135.21, df = 2 | p < 0.001 |
| **Emotion Regulation** | 64.96, 18.27<sup>a</sup> | 79.13, 20.93<sup>b</sup> | 96.82, 23.59<sup>c</sup> | F = 151.46, df = 2 | p < 0.001 |
| **Anxiety** | 8.82, 9.32<sup>a</sup> | 14.93, 10.29<sup>b</sup> | 23.43, 13.14<sup>c</sup> | F = 112.5, df = 2 | p < 0.001 |
| **Depression** | 7.41, 8.28<sup>a</sup> | 13.66, 10.61<sup>b</sup> | 22.74, 12.72<sup>c</sup> | F = 133.05, df = 2 | p < 0.001 |
| **Impulsivity** | 54.22, 9.52<sup>a</sup> | 59.91, 9.50<sup>b</sup> | 65.60, 11.37<sup>c</sup> | F = 81.20, df = 2 | p < 0.001 |
| **Positive Affect** | 30.47, 9.32<sup>a</sup> | 27.47, 7.70<sup>b</sup> | 23.89, 8.28<sup>c</sup> | F = 39.25, df = 2 | p < 0.001 |
| **Negative Affect** | 11.61, 3.68<sup>a</sup> | 14.05, 5.90<sup>b</sup> | 17.32, 7.51<sup>c</sup> | F = 60.92, df = 2 | p < 0.001 |
| **Physical QOL** | 16.07, 2.65<sup>a</sup> | 14.94, 2.74<sup>b</sup> | 13.58, 3.11<sup>c</sup> | F = 41.47, df = 2 | p < 0.001 |
| **Psychol. QOL** | 15.04, 2.71<sup>a</sup> | 13.24, 2.89<sup>b</sup> | 11.04, 2.90<sup>c</sup> | F = 102.62, df = 2 | p < 0.001 |
| **Social QOL** | 14.83, 3.30<sup>a</sup> | 13.14, 3.69<sup>b</sup> | 11.86, 3.77<sup>c</sup> | F = 47.51, df = 2 | p < 0.001 |
| **Environ. QOL** | 15.26, 2.47<sup>a</sup> | 14.06, 2.61<sup>b</sup> | 12.92, 2.61<sup>c</sup> | F = 46.74, df = 2 | p < 0.001 |
| **Neuroticism** | 27.80, 8.16<sup>a</sup> | 34.69, 7.30<sup>b</sup> | 40.44, 7.65<sup>c</sup> | F = 717.86, df = 2 | p < 0.001 |
| **Extraversion** | 41.59, 8.34<sup>a</sup> | 39.21, 8.46<sup>b</sup> | 37.54, 8.62<sup>c</sup> | F = 62.43, df = 2 | p < 0.001 |
| **Openness** | 40.92, 7.27<sup>a</sup> | 42.80, 7.15<sup>b</sup> | 43.79, 6.96<sup>c</sup> | F = 45.16, df = 2 | p < 0.001 |
| **Agreeableness** | 49.12, 5.92<sup>a</sup> | 47.74, 5.97<sup>b</sup> | 46.12, 6.56<sup>c</sup> | F = 63.72, df = 2 | p < 0.001 |
| **Conscientiousness** | 50.83, 6.12<sup>a</sup> | 47.58, 6.41<sup>b</sup> | 44.04, 6.87<sup>c</sup> | F = 296.36, df = 2 | p < 0.001 |

*Note.* Post hoc test for ANOVAs is Bonferroni, z-test was used to compare column proportions after Chi-square. Superscript letters that differ across comparison groups indicate statistically significant difference, $p < .05$ . Any groups that have the same letters did not statistically differ from each other. Scales (see methods for details): BPD symptoms – BSL23, Emotion Regulation – DERS, Anxiety – BAI, Depression – BDI-II, Impulsivity – BIS, Positive and Negative Affect – PANAS, 4 QOL subscales – WHOQOL, Big 5 – IPIP60.

We tested our participants’ endorsement of BPD symptoms for reliability by measuring the correlation between score on the BPD symptom report in HIT1 (SCID-II self-report questionnaire) and BPD symptom report in HIT2 (BSL-23), which occurred days to weeks later (mean = 16.8 days). The number of self-reported symptoms of BPD on the SCID-II and the total score on the BSL-23 were positively correlated at *r* = .58, *p <* .001.

### Psychiatric co-morbidities by group

In HIT2, we examined symptoms and personality traits of participants in each group. (**Table 3: HIT2 Scales**). We found the expected significant between-group differences, with Current BPD > Remitted BPD > Never BPD on all scales (post-hoc tests all *p <* 0.05). Specifically, people with BPD endorsed more BPD symptoms (*F*(2, 3018) = 135.21, *p 003C* 0.001, η^2^ = 0.295), more difficulties with emotion regulation (*F*(2, 550) = 151.46, *p* < 0.001, η^2^ = 0.301), more anxiety (*F*(2, 493) = 112.5, *p* < 0.001, η^2^ = 0.272), more depression (*F*(2, 537) = 133.05, *p* < 0.001, η^2^ = 0.297), more impulsivity (*F*(2, 476) = 81.20, *p* < 0.001, η^2^ = 0.207), and more negative affect (*F*(2, 543) = 60.92, *p* < 0.001, η^2^ = 0.134) than did people without BPD. Also, people with BPD endorsed less positive affect (*F*(2, 545) = 39.25, *p* < 0.001, η^2^ = 0.074) and lower quality of life (subscales with F-tests ranging from *F*(2, 550) = 41.47 – 102.62, *p* < 0.001, η^2^ = 0.131 – 0.272) than did people without BPD.

We also observed the expected correlations between symptom scale scores (**Table S1**).

### Personality traits of participants with current, remitted, and never BPD

Participants with BPD also reported personality traits consistent with past reports (**Table 3**). These data, collected in HIT1, included the full sample of 3,021 individuals, and we found significant differences (post-hoc tests p < 0.001) between each of the three groups for each of the big five traits (Current BPD vs. Remitted BPD vs. Never BPD). People with BPD endorsed higher trait neuroticism, *F*(2, 3018) = 717.86, *p* < 0.001, η^2^ = 0.322, and openness, *F*(2, 3018)= 45.16, *p* < 0.001, η^2^ = 0.029, and lower extraversion, *F*(2, 3018) = 63.43, *p* < 0.001, η^2^ = 0.040, agreeableness, *F*(2, 3018) = 63.72, *p* < 0.001, η^2^ = 0.041, and conscientiousness, *F*(2, 3018) = 296.36, *p* < 0.001, η^2^ = 0.164.

### History of self-harm in participants with current, remitted, and never BPD

An examination of the chronology of self-harm in each group revealed significant and large between-group differences (**Figure 1,** tested 3 groups x 3 time frames (lifetime, past, in the last month): X^2^ (4) = 602.34, *p* < 0.001, Cramer’s V = 0.316. Lifetime history of self-harm was more frequent in people with BPD (Never BPD 4.7%, Remitted BPD 27.3%, Current BPD 50%, and post-hocs revealed significant differences between each of the 3 groups (*p* < 0.05; **Figure 1A**). Past history of self-harm was also more frequent in people with BPD (Never BPD 4.6%, Remitted BPD 27.1%, Current BPD 44.6%, and post-hoc tests revealed significant differences between each of the 3 groups (*p* < 0.05; **Figure 1B**). Finally, past month history of self-harm was also more frequent in people with BPD (Never BPD 0.10%, Remitted BPD 0.20%, Current BPD 5.50%, however, here post-hocs distinguished only the Current BPD group from the other two groups (*p* < 0.05; **Figure 1C**).

**Figure 1.**
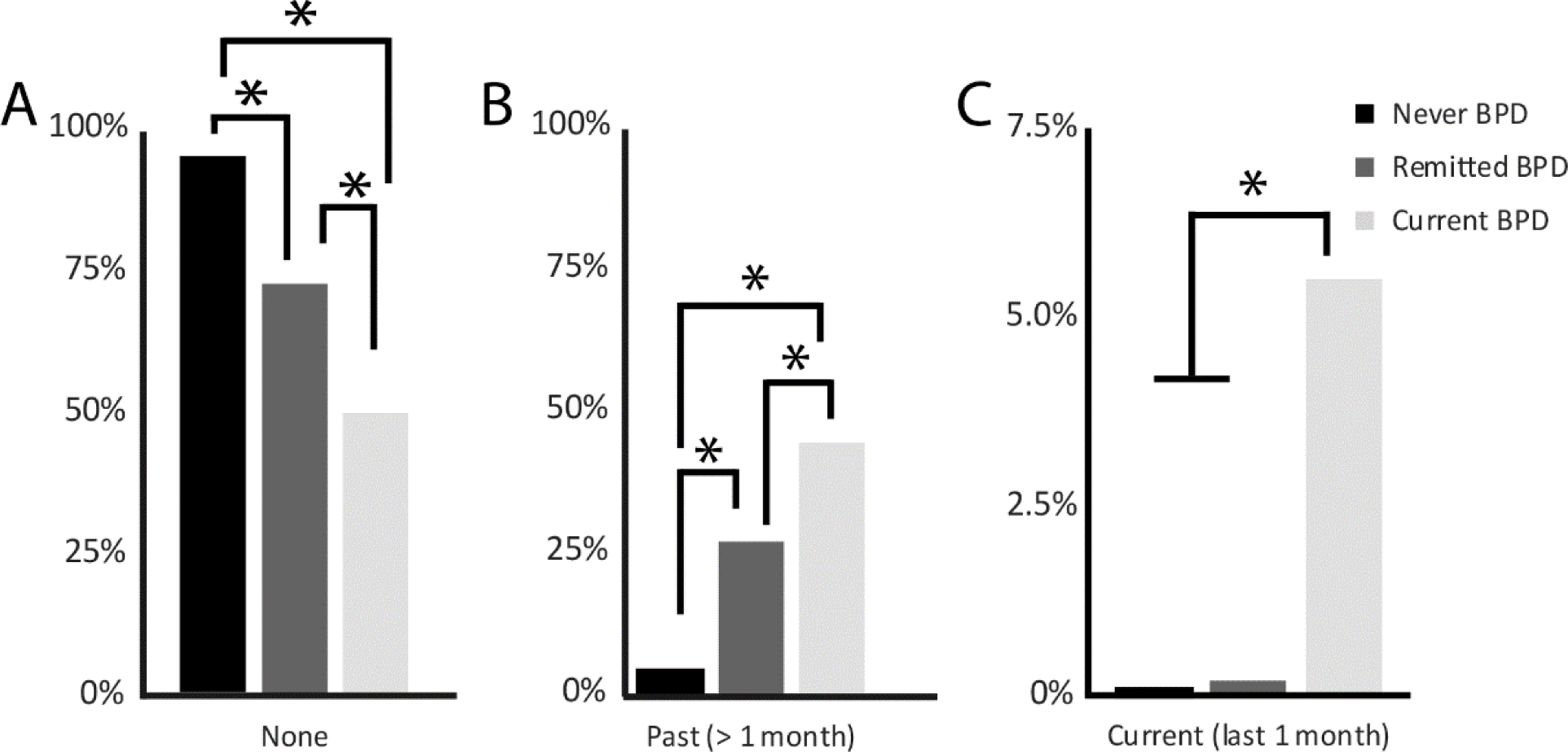
Self harm. Frequency of participants reporting A) no history of self harm, B) past self-harm, and C) self-harm in the past month is displayed by group: Never BPD (black bars), Remitted BPD (dark gray bars), Current BPD (light grey bars). * denotes p < 0.05.

## Discussion

In this study, we aimed to test MTurk as a potential venue for BPD research. We compared prevalence and associated factors in a large MTurk sample to previously established data from community and patient samples. We tested for respondents’ careful completion of surveys and reliability of their responses. We measured demographics and frequency of past mental health diagnosis and of self-reported personality traits and BPD symptoms now and in the past. We also measured these features in the Remitted BPD group. Overall, we found that BPD features are quite frequent among MTurk workers, both in terms of direct interrogation of DSM criteria, and expected features of social life, symptoms, and personality.

### Overall sample

Our sample was similar to other researchers’ MTurk samples in terms of demographics and self-reported psychopathology (Berinsky et al., 2012; Buhrmester et al., 2011; Paolacci &Chandler, 2014; Shapiro, Chandler, & Mueller, 2013). With regard to BPD symptoms in particular, we found a very high frequency of self-reported symptoms, with one-third of our participants meeting DSM criteria for BPD. This is far higher than other large community samples that have found prevalence rates to be closer to 5-6% (Grant et al., 2008).

We tested for consistency and reliability of responding, and found high correlations between scores on our two different BPD self-reports, and between BPD scores and mood and emotional regulation symptom scales. We also found that BPD scores correlated with lower self-reported quality of life, lower positive affect, higher negative affect, and higher neuroticism. Meeting criteria for BPD in our sample also predicted more psychiatric medication use and higher rates of obesity and high blood pressure. These results are consistent with the expected profile for people with BPD.

There are several possible explanations for this surprisingly high frequency of people with BPD in our sample, in addition to the higher base rates typically yielded by self-report measures (Hopwood et al., 2008). MTurk may be attractive to people with BPD, as it allows for social interactions with boundaries set by task directions, durations, and contracts, but also allows the MTurk worker a great deal of flexibility in aspects of engagement such as where to work, when to work, how long to work, what to work on, and who to work with. This combination of clear social structure and personal flexibility may facilitate participation by this group of people with high anxiety, fluctuating symptoms, and social difficulties.

Another consideration is whether our symptom measures are specific tests for BPD or more general markers of psychopathology. We did observe high correlations between BPD, depression, and anxiety self-report measures, consistent with known patterns of co-morbidity in BPD. We also note that in our sample, among people who met criteria for current BPD, 50% had at some point in their lives tried to or threatened to hurt or kill themselves, consistent with chronic suicidality and marked increases in the frequency of suicidal action in BPD. Additionally, although self-report methods of obtaining diagnoses cannot replace a diagnosis given by a clinician following a structured diagnostic interview, the SCID-II-PQ has been found to be sensitive and specific, with moderate-to-excellent accuracy when compared to clinical interviews (Fowler et al., 2018).

### Discrepancy between high frequency of meeting BPD criteria and low frequency of past diagnosis

Only a small number of those meeting criteria for BPD reported a previous <u>diagnosis</u> of BPD, though many carried diagnoses of mood and anxiety disorders. The discrepancy between symptom-reports and previous diagnosis may arise because of under-diagnosis or misdiagnosis in the community. This is a frequent problem, likely driven by lack of clinician experience in BPD diagnosis, or preference for giving a diagnosis with less stigma that is perceived to be more trea316356 (e.g. (Gunderson, 2009).

### Remission

Our approach to defining a remitted BPD group here depends on the participant’s retrospective recall of their past symptoms. This is a major limitation of this approach. However, report of current symptoms and social status cohere with the claim that people in this group are at an intermediate level of difficulty. We hope that future work on BPD in this setting will probe remission and recovery in more detail to better understand this group. We think that both the Current and Remitted BPD groups here likely include individuals who, on detailed clinical interview, may be more in the BPD traits (< 5 DSM criteria) range than the full syndrome range. However, some people may also under-report symptoms to a novel interviewer but feel comfor316356 in the potentially more anonymous feeling setting of an online survey. Identifying how life experience changes as BPD symptoms are ameliorated, and what factors may predict and promote recovery, are critical questions. The MTurk worker population does appear to include a large number of people with high and mid-range levels of BPD symptoms.

### Potential implications of findings

We have defined the frequency and characteristics of MTurk workers who meet criteria for current or remitted BPD. Our findings suggest that MTurk will be a good locus for future work to test survey response and potentially also cognitive tasks in people with BPD. Large sample sizes in this setting hold promise for doing studies that provide more statistical power, better engage a dimensional approach, and that consider the experiences of people in remission and in recovery.

---

**Table S1.** Correlations between measures administered in study phase two.

|  | BSL | DERS | BAI | BDI | BIS | QOL<br>Health | QOL<br>Psych | QOL<br>SR | QOL<br>Environ | PANAS<br>Pos | PANAS<br>Neg |
| --- | --- | --- | --- | --- | --- | --- | --- | --- | --- | --- | --- |
| BSL | - |  |  |  |  |  |  |  |  |  |  |
| DERS | **.68** | - |  |  |  |  |  |  |  |  |  |
| BAI | **.75** | **.54** | - |  |  |  |  |  |  |  |  |
| BDI | **.82** | **.72** | **.65** | - |  |  |  |  |  |  |  |
| BIS | **.49** | **.62** | **.42** | **.54** | - |  |  |  |  |  |  |
| QOL Health | **-.54** | **-.46** | **-.56** | **-.61** | **-.42** | - |  |  |  |  |  |
| QOL Psych. | **-.65** | **-.65** | **-.51** | **-.76** | **-.49** | **.65** | - |  |  |  |  |
| QOL SR | **-.39** | **-.41** | **-.31** | **-.51** | **-.33** | **.48** | **.62** | - |  |  |  |
| QOL Envir. | **-.44** | **-.43** | **-.40** | **-.53** | **-.33** | **.63** | **.65** | **.55** | - |  |  |
| PANAS Pos | **-.27** | **-.42** | **-.19** | **-.36** | **-.32** | **.28** | **.56** | **.40** | **.34** | - |  |
| PANAS Neg | **.66** | **.57** | **.52** | **.61** | **.45** | **-.39** | **-.41** | **-.27** | **-.32** | **-.14** | - |

*Note:* bolded values are statistically significant ( $p < .01$ ).

